# A network of microRNAs acts to promote cell cycle exit and differentiation of human pancreatic endocrine cells

**DOI:** 10.1101/618330

**Authors:** Wen Jin, Francesca Mulas, Bjoern Gaertner, Yinghui Sui, Jinzhao Wang, Chun Zeng, Nicholas Vinckier, Allen Wang, Kim-Vy Nguyen-Ngoc, Joshua Chiou, Klaus H. Kaestner, Kelly Frazer, Andrea C. Carrano, Hung-Ping Shih, Maike Sander

## Abstract

Pancreatic endocrine cell differentiation is orchestrated by transcription factors that operate in a gene regulatory network to activate endocrine lineage genes and repress lineage-inappropriate genes. MicroRNAs (miRNAs) are important modulators of gene expression, yet their role in endocrine cell differentiation has not been explored system-wide. Here we characterize miRNA-regulatory networks active in human endocrine cell differentiation by combining small RNA sequencing, miRNA overexpression experiments, and network modeling approaches. This analysis identifies Let-7g, Let-7a, miR-200a, and miR-375 as endocrine-enriched miRNAs with high impact on driving endocrine differentiation-associated gene expression changes. These miRNAs target different sets of transcription factors, which converge on a network of genes involved in cell cycle regulation. When expressed in human embryonic stem cell-derived pancreatic progenitors these miRNAs induce cell cycle exit and promote endocrine cell differentiation. Our study delineates the role of miRNAs in human endocrine cell differentiation and identifies miRNAs that could facilitate endocrine cell reprogramming.

## INTRODUCTION

The potential to generate pancreatic beta cells from human pluripotent stem cells (hPSCs) or via cell reprogramming from other cell sources holds promise for modeling causes of diabetes and cell replacement therapies (Benthuysen et al., 2016). Knowledge of the molecular underpinnings of pancreas and beta cell differentiation has enabled some success in developing beta cell reprogramming and directed differentiation strategies. In particular, the identification of transcription factors (TFs) governing cell fate decisions has been instrumental for cell reprogramming approaches (Benthuysen et al., 2016). While TFs play a major role in orchestrating gene expression changes during developmental transitions, recent evidence also shows significant roles for other regulators such as small RNAs.

MicroRNAs (miRNAs) are a group of small non-coding RNAs (∼22 nucleotides) with known roles in the regulation of gene expression in development, mature cell function, and disease (Vidigal and Ventura, 2015). Studies in mice and zebrafish have demonstrated important roles for miRNAs in pancreatic endocrine cell development and beta cell function (Kaspi et al., 2014). Pancreatic progenitor cell-specific deletion of *Dicer1,* an enzyme that is universally required for the functional maturation of miRNAs, results in reduced endocrine cell numbers (Lynn et al., 2007), while *Dicer1* disruption in beta cells impairs insulin biogenesis (Melkman-Zehavi et al., 2011). At the level of individual miRNAs, beta cell-enriched miR-375 (Kloosterman et al., 2007; Poy et al., 2009) and miR-7 (Kredo-Russo et al., 2012; Latreille et al., 2014) have been identified as regulators of beta cell differentiation and function.

miRNAs functionally repress target mRNAs and act predominately by destabilizing mRNAs through base pairing between the miRNA seed sequence (nucleotides at position 2–8) and a complementary sequence in the target mRNA (Guo et al., 2010; Lim et al., 2005). The effects of individual miRNAs on gene expression are generally small, which has led to the concept that miRNAs fine-tune gene expression rather than acting as genetic switches (Vidigal and Ventura, 2015). Consistent with this idea, miRNAs have been shown to promote cell differentiation and to facilitate cell reprogramming when force expressed in conjunction with lineage-determining TFs (Chen et al., 2004; Chen et al., 2006; Dey et al., 2012; Lim et al., 2005; Nam et al., 2013; Yoo et al., 2011). Mechanistically, each miRNA has the ability to repress hundreds of mRNA targets, and multiple miRNAs often converge on a single pathway to promote a common developmental outcome (Lim et al., 2005; Vidigal and Ventura, 2015). Therefore, a comprehensive understanding of context-specific contributions of miRNAs to gene regulation requires a systems-level approach where all miRNAs and their targets are considered.

In this study we used genome-wide small RNA sequencing to identify candidate miRNAs with possible roles in human beta cell differentiation. By comparing miRNA profiles of hPSC-derived pancreatic progenitors and human cadaveric beta cells genome-wide, we identified miRNAs that are induced during beta cell differentiation. Through gain-of-function experiments during hPSC differentiation, we show that beta cell-enriched miRNAs act synergistically to promote cell cycle exit and beta cell differentiation. Integrating RNA-seq, CLIP-seq and chromatin state data, we applied a network modeling approach to identify specific miRNA-regulated TFs that explain the impact of beta cell-enriched miRNAs on cell cycle regulation during beta cell differentiation. Our findings provide a systems level view of how miRNAs regulate human beta cell differentiation, which has implications for programming beta cells from hPSCs or other cell sources.

## RESULTS

### Identification of miRNAs upregulated during endocrine cell differentiation

To identify miRNAs that are regulated during pancreatic beta cell differentiation, we conducted genome-wide small RNA sequencing in pancreatic progenitor cells derived from CyT49 human embryonic stem cells (hESCs) (**Figure S1**) and primary beta cells isolated from cadaveric human islets by fluorescence activated cell sorting (Kameswaran et al., 2014) (**Figure 1A**). By comparing expression levels of individual miRNAs in beta cells and pancreatic endoderm stage (PE) cells, we defined miRNAs induced during beta cell differentiation. This analysis revealed 13 miRNAs that were more highly expressed in beta cells than in PE cells (> 5000 sequence reads in beta cells; > 2-fold increase; **Figure 1B**; **Table S1A,B**). With the exception of miR-127, miR-204, and miR-99b, the same miRNAs also exhibited higher expression in sorted alpha cells compared to PE cells (**Figure 1C**; **Table S1A,C**), suggesting shared roles for most miRNAs in the development of both endocrine cell types. Among the miRNAs induced during endocrine cell differentiation were miR-375, miR-200a/c, and miR-7, which have reported roles in beta cell development, beta cell proliferation, function, and survival in mice (Belgardt et al., 2015; Kloosterman et al., 2007; Kredo-Russo et al., 2012; Latreille et al., 2014; Nieto et al., 2012; Poy et al., 2004; Poy et al., 2009; Wang et al., 2013). Most notable was the significantly higher expression of members of the Let-7 miRNA family in both beta and alpha cells compared to PE cells, including Let-7a, Let-7b, Let-7f, Let-7g, and miR-98 (**Figure 1B,D**; **Table S1A,B**). We confirmed the results from the small RNA sequencing by comparing miRNA levels in PE cells and human cadaveric islets using the Taqman miRNA assay (**Figure 1E**).

**Figure 1.**
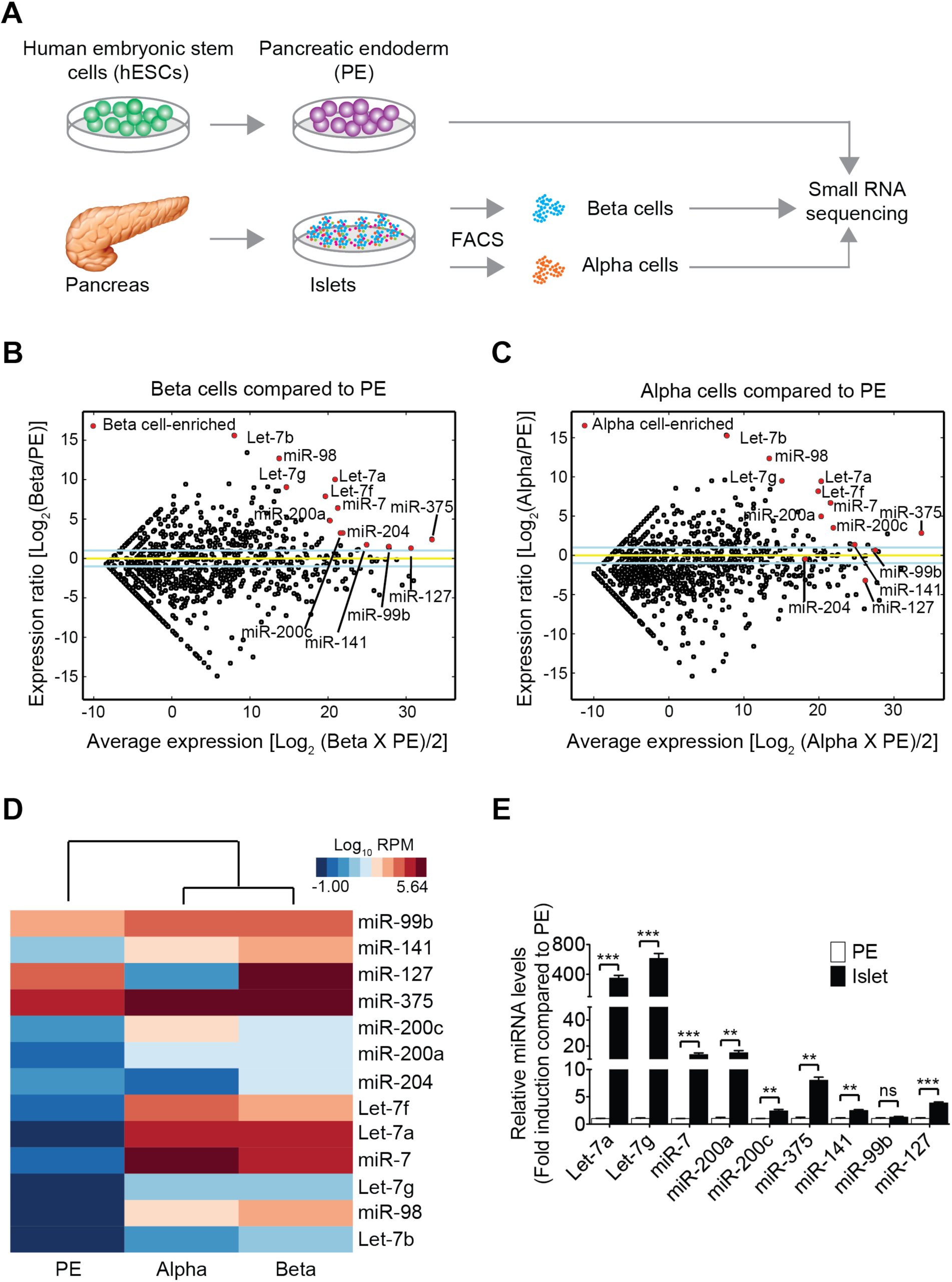
Identification of miRNAs up-regulated during endocrine cell differentiation of human pancreatic progenitor cells. **(A)** Workflow for genome-wide small RNA profiling of pancreatic progenitors (pancreatic endoderm, PE) and endocrine islet cells. PE cells were differentiated from human embryonic stem cells (hESCs) and human alpha and beta cells were isolated from cadaveric human islets by fluorescence activated cell sorting (FACS). **(B,C)** MA plots comparing miRNA expression levels in PE cells and beta cells **(B)** or PE cells and alpha cells **(C)**. miRNAs with higher expression in beta cells than PE are indicated by red circles in **B** and **C**. Blue lines indicate two-fold change in miRNA expression, yellow line indicates no change. **(D)** Heatmap comparing expression levels in PE, alpha cells and beta cells of the thirteen most highly enriched miRNAs in beta cells compared to PE cells. **(E)** Relative expression of indicated miRNAs determined by Taqman qPCR in PE cells and human islets. Data are shown as mean ± S.E.M. (n = 3 biological replicates). ns, not significant; \*\**p* < 0.01, \*\*\**p* < 0.001; Student’s t-test. See also Figure S1 and Table S1.

### Identifying candidate miRNAs regulating human endocrine cell differentiation

To identify mRNAs regulated by these miRNAs, we next chose the eight miRNAs most highly regulated between PE and beta cells for over-expression experiments in HeLa cells. All Let-7 family members share the same seed sequence, which is important for target recognition, and therefore are predicted to regulate similar targets. Therefore, of the numerous Let-7 family members enriched in beta cells (**Figure 1B**; **Table S1A,B**), we over-expressed only Let-7a and Let-7g, the two Let-7 miRNAs most differentially regulated between PE and beta cells. Since miR-141 and miR-200c also share a common seed sequence, we force-expressed only miR-200c which exhibited higher induction from PE to islet than miR-141. As previous studies have shown that cellular context does not significantly affect miRNA target repertoires (Nam et al., 2014), we used HeLa cells to transfect individual precursor miRNAs and harvested cells 24 hours later for RNA analysis (**Figure 2A**). Quantitative PCR analysis demonstrated that individual mature miRNAs were efficiently induced in HeLa cells (**Figure S2A**). Furthermore, we observed reduced expression of their published target mRNAs, confirming biologically relevant expression levels of the miRNAs (Latreille et al., 2015; Liu et al., 2015; Mayr et al., 2007; Park et al., 2008; Yu et al., 2013) (**Fig. S2B**). We next identified coding genes repressed by over-expression of each miRNA (*p* < 0.05, permutation test; **Table S2A-I**) and determined whether miRNA-repressed mRNAs are also down-regulated during the transition from PE to islet cells. We used Gene Set Enrichment Analysis (GSEA) (Subramanian et al., 2005) to score the a-priori defined sets of miRNA-repressed mRNAs for expression differences between PE and islet cells. mRNAs repressed by Let-7g, Let-7a, miR-200a, and miR-375 exhibited significantly lower transcript levels in islets than in PE cells (**Figure 2B**). By contrast, mRNAs repressed by miR-99b, miR-127, miR-7, and miR200c either exhibited significantly lower levels in PE cells than in islet cells (*p* = 0.022 for miR-99b; *p* = 0.027 for miR-7) or no significant difference between PE and islet cells (*p* = 0.57 for miR-127; *p* = 0.147 for miR200c). Together, these results indicate that out of the initial eight miRNA candidates, Let-7g, Let-7a, miR-200a, and miR-375 have the highest predictive value of regulating endocrine cell differentiation.

**Figure 2.**
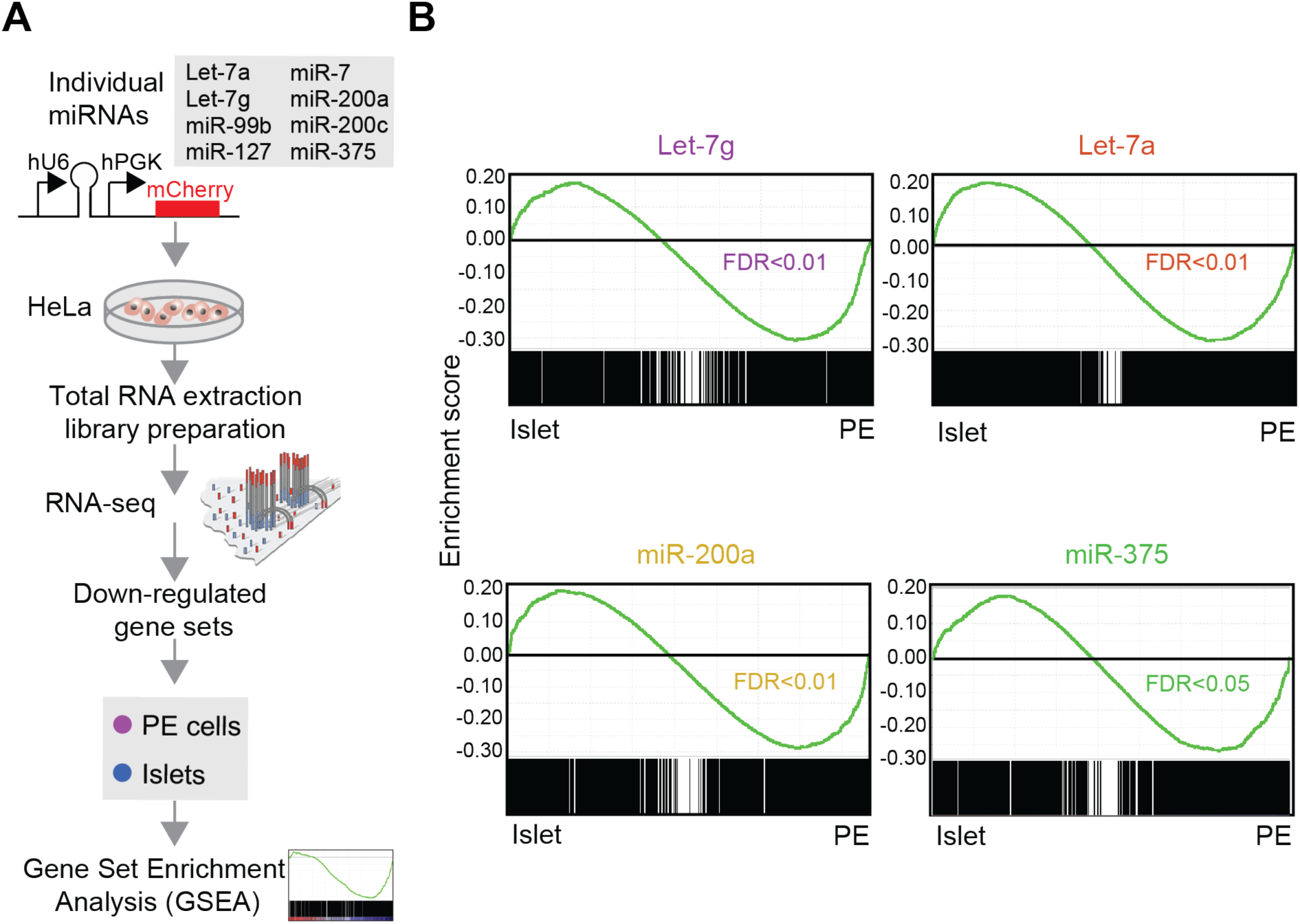
Identification of beta cell-enriched miRNAs most predictive of regulating human endocrine cell differentiation. **(A)** Workflow to identify genes repressed by each indicated miRNA after transfection of miRNA-expressing plasmids into HeLa cells and RNA-seq analysis 24 h later (n = 1). Down-regulated genes were subjected to Gene Set Enrichment Analysis (GSEA). **(B)** GSEA plots showing enrichment of genes repressed by Let-7g, Let-7a, miR-200a, and miR-375 in islets (n = 3 donors) compared to PE (n = 2 biological replicates). False Discovery Rate (FDR) is shown. See also Figure S2 and Table S2.

To demonstrate that findings in HeLa cells are relevant in pancreatic progenitor cells, we over-expressed Let-7g, Let-7a, miR-200a, and miR-375 individually in hESC-derived PE cells (**Figure 3A**). For these studies, we chose PE cells derived from H1 hESCs because a recently published protocol showed very efficient differentiation of H1 hESCs into beta-like cells in vitro (Rezania et al., 2014). Since our genome-wide small RNA sequencing was performed in PE cells from CyT49 hESCs (**Figure 1**), we first confirmed that H1 and CyT49 hESC-derived PE cells have similar molecular features. Similar to CyT49 hESC-derived PE cells (**Figure S1**), 98% H1 hESC-derived PE cells expressed the pancreatic progenitor marker PDX1 (**Figure S3A,B**). In addition, RNA-seq analysis showed highly concordant transcriptome profiles of H1 and CyT49 hESC-derived PE cells [(R) > 0.92; **Figure S3C**]. Furthermore, we confirmed that Let-7g, Let-7a, miR-200a, and miR-375 were expressed at similarly low levels in H1 and CyT49 hESC-derived PE cells (**Figure S3D**).

**Figure 3.**
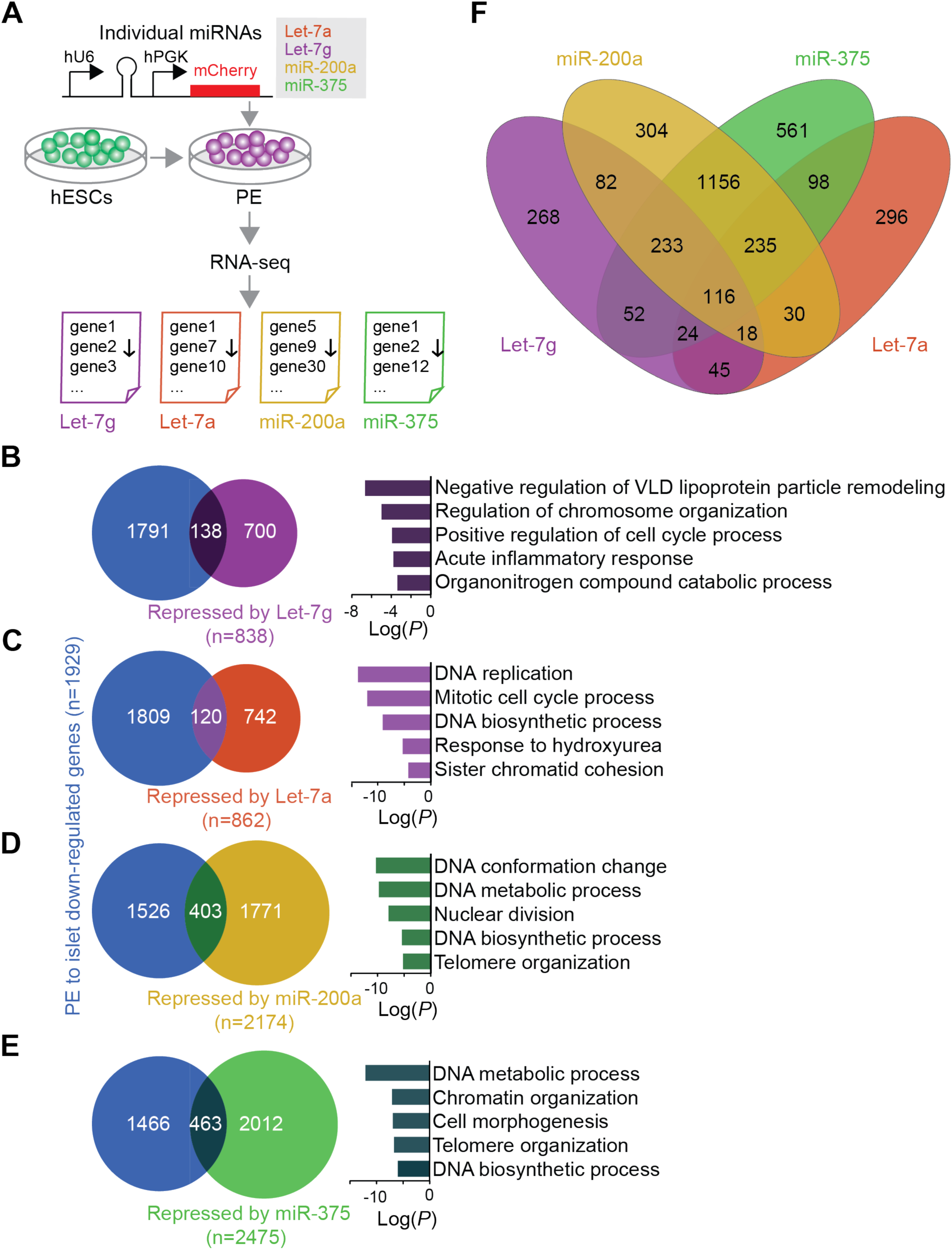
Beta cell-enriched miRNAs regulate expression of cell cycle genes in pancreatic progenitor cells. **(A)** Workflow to identify genes repressed by each indicated miRNA after lentiviral transduction of hESC-derived pancreatic endoderm (PE) cells. Transduced cells were sorted based on mCherry after 48 h, RNA-seq analysis performed (n = 3 biological replicates), and down-regulated genes identified. **(B-E)** Venn diagrams showing the overlap between genes down-regulated in islets (n = 3) compared to PE (n = 2) (blue) and genes repressed by Let-7g (purple, **B**), Let-7a (red, **C**), miR-200a (yellow, **D**), or miR-375 (green, **E**). Top five GO categories enriched among genes repressed by the miRNA and down-regulated in islets compared to PE are shown on the right. **(F)** Venn diagram showing overlap between miRNA-repressed genes. See also Figure S3, Table S3, and Table S4.

Inclusion of a mCherry reporter into the miRNA constructs allowed us to monitor transduction efficiencies in PE stage cultures and to isolate transduced cells by FACS. We observed 13-20% mCherry^+^ PE cells two days after transduction, and this number increased to 34-49% by day six (**Figure S3E**). The increase is likely explained by the lentiviral expression vector requiring more than two days to reach maximum expression. To identify miRNA targets, we chose to analyze sorted mCherry^+^ PE cells two days after transduction, reasoning that this early time point is best suited for studying the direct effects of miRNA on gene expression. As expected, Let-7g, Let-7a, miR-200a, and miR-375 were each significantly higher expressed in cells transduced with the miRNA-expressing vector compared to control vector-transduced cells (**Figure S3F**). Furthermore, as in HeLa cells (**Figure 2B**), forced expression of Let-7g, Let-7a, miR-200a, or miR-375 in hESC-derived PE repressed the expression of genes (*p* < 0.05, permutation test, **Table S3A-D**) that were down-regulated between PE and islets (**Figure S3G**), suggesting that these miRNAs could contribute to gene expression changes during islet cell differentiation.

### Beta cell-enriched miRNAs regulate expression of cell cycle genes in pancreatic progenitor cells

To identify miRNA-regulated transcripts with likely roles in endocrine cell differentiation, we analyzed sets of genes that were down-regulated by forced expression of each miRNA and also down-regulated in islets as compared to PE cells (*p* < 0.05, permutation test, **Figure 3B-E**). These mRNA subsets comprised 16.5% of Let-7g-, 13.9% of Let-7a-, 18.5% of miR-200a-, and 18.7% of miR-375-repressed mRNAs in PE cells. We then performed Gene Ontology (GO) analysis to define the biological processes regulated by mRNAs that are repressed by individual miRNAs and are also expressed at lower level in islet than PE cells. The top five enriched GO categories for each one of these miRNA-regulated sets of mRNAs comprised processes associated with DNA replication and regulation of the cell cycle (**Figure 3B-E**; **Table S4A-D**). Given that endocrine cell formation is associated with cell cycle exit (Kim et al., 2015; Miyatsuka et al., 2011; Piccand et al., 2014), these findings suggest that miRNAs could control endocrine cell differentiation by regulating mRNAs involved in cell cycle control. The finding that all four miRNAs regulate cell cycle-associated transcripts raised the question of whether they share similar targets. Analysis of the extent of overlap between the mRNAs down-regulated by Let-7g, Let-7a, miR-200a, and miR-375 revealed a modest number of shared targets (**Figure 3F**). Only 116 mRNAs were repressed by all four miRNAs, indicating distinct regulatory roles for each one of the four miRNAs. Together, these results suggest distinct but converging miRNA targets in regulating cell division in pancreatic progenitors.

Since all four candidate miRNAs appeared to regulate different aspects of cell cycle progression, we sought to gain further insight into how input from the different miRNAs converges on cell cycle regulation. To study the combined effect of all four miRNAs, we generated a “polycistronic” miRNA (poly-miR) lentiviral construct that drives the expression of all four miRNAs under the control of a single promoter. We expressed the poly-miR construct in H1 hESC-derived PE cells and analyzed the transcriptome two days after transduction (**Figure 4A**). miRNA expression analysis in mCherry-sorted cells revealed that Let-7g, Let-7a, miR-200a, and miR-375 were each significantly higher expressed in poly-miR- than vector-only-transduced PE cells (**Figure S4A,B**). Expression of the poly-miR construct in PE cells resulted in down-regulation of 2,463 transcripts (*p* < 0.05; permutation test). Consistent with the results from expression of individual miRNAs (**Figure S3G**), poly-miR-repressed mRNAs (*p* < 0.05, permutation test, **Table S3E**) were highly enriched for mRNAs with higher expression in PE compared to islets (**Figure S4C**). Of the 2,463 poly-miR-repressed mRNAs, 388 were also down-regulated during the transition of PE to islet (**Figure S4D**). As predicted, genes involved in cell cycle processes were overrepresented among these 388 mRNAs (**Figure S4D**; **Table S4E**).

**Figure 4.**
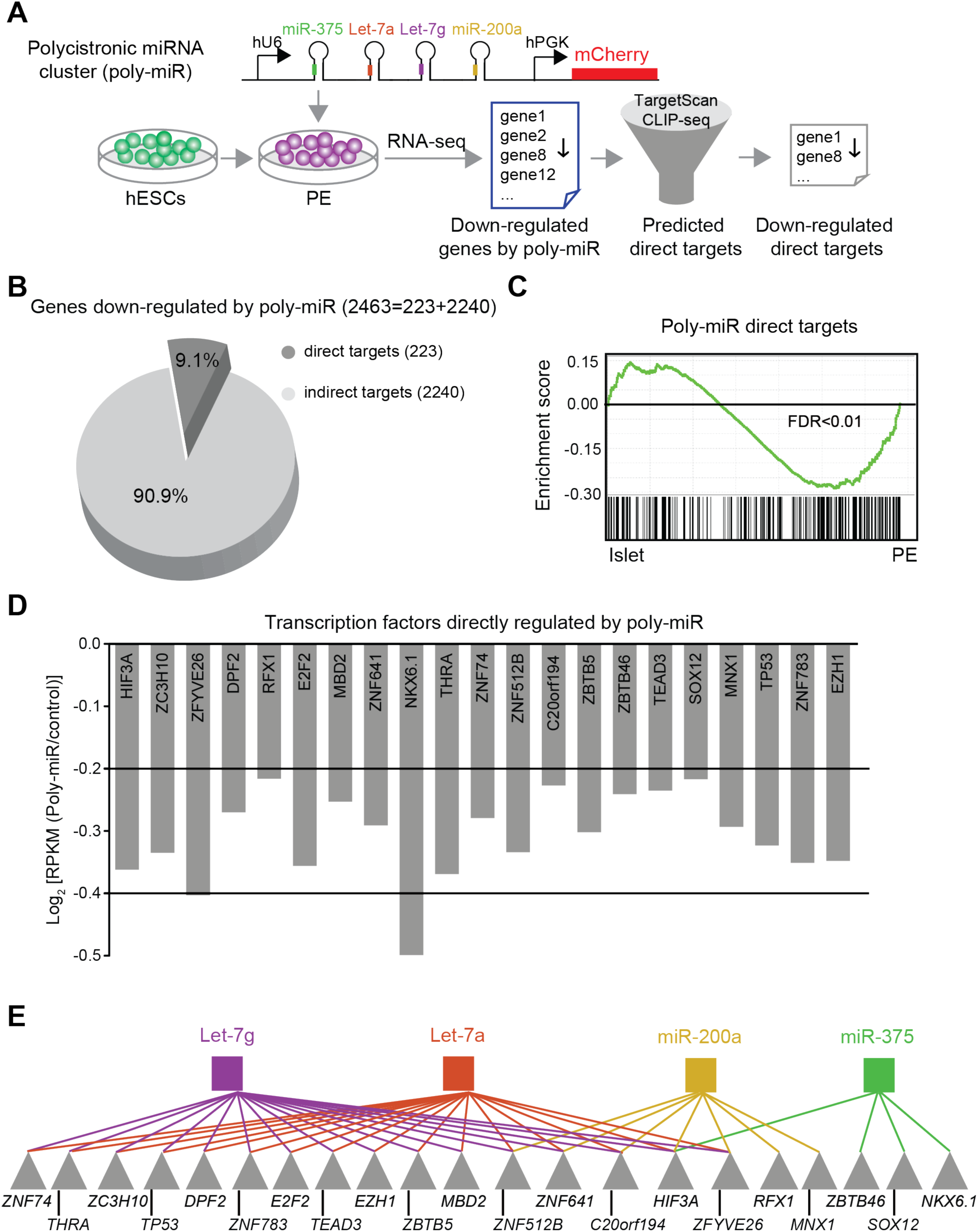
Identification of direct miRNA target mRNAs in pancreatic progenitor cells. **(A)** Workflow to identify repressed genes after transduction of hESC-derived pancreatic endoderm (PE) cells with a lentivirus expressing a polycistronic construct for the indicated miRNAs (poly-miR) and mCherry. Transduced cells were sorted after 48 h, RNA-seq analysis performed (n = 3 biological replicates), and down-regulated genes identified. Direct targets of candidate miRNAs were identified based on TargetScan and CLIP-seq analysis. **(B)** Pie graph showing percentage of direct (dark grey) and indirect (light grey) targets of candidate miRNAs repressed by poly-miR construct. **(C)** GSEA plot showing enrichment of 223 direct target genes of Let-7g, Let-7a, miR-200a, and miR-375 in islets (n = 3) compared to PE (n = 2). False Discovery Rate (FDR) is shown. **(D)** mRNA expression levels of transcription factors predicted to be directly targeted by Let-7g, Let-7a, miR-200a, and miR-375 measured in reads per kilobase per million reads mapped (RPKM). **(E)** Predicted network of transcription factors downstream of miRNAs. Transcription factors are indicated by grey triangles and individual miRNAs are indicated by colored squares. See also Figure S4, Table S3, Table S4, and Table S5.

### Cell-cycle-associated transcription factors are predicted direct targets of beta cell-enriched miRNAs

To decipher mechanisms by which Let-7g, Let-7a, miR-200a, and miR-375 regulate cell cycle genes, we sought to distinguish direct and indirect targets of the four miRNAs (**Figure 4A**). We defined putative direct targets as poly-miR-repressed mRNAs (*p* < 0.05, permutation test, **Table S3E**) predicted to be direct targets by TargetScan (based on matching sequence to the miRNA seed region) and/or exhibiting binding to the RNA-binding protein Argonaute, as determined by CLIP-seq in human islets (Kameswaran et al., 2014). From this analysis, 223 direct target mRNAs, which represent 9.1% of all poly-miR-repressed genes, were identified (**Figure 4B**; **Table S5A**). Reinforcing the potential relevance of these direct miRNA targets for endocrine cell development, GSEA analysis showed significantly lower expression of these genes in islets than in PE cells (**Figure 4C**).

To determine whether miRNAs are direct regulators of cell cycle-associated mRNAs in PE cells, we analyzed enriched GO terms among the 223 direct miRNA targets. We found no enrichment of categories linked to cell cycle-related processes (**Table S5B**). Moreover, many of the cell cycle regulators that were repressed by the poly-miR, including *CCND3*, *CDC45*, *MCM7*, and *CKS1B* (**Table S3E**), were not among the direct miRNA targets (**Table S5A**). Thus, cell cycle-associated transcripts appear to be indirectly regulated by the miRNAs. We postulated that this indirect effect of miRNAs on the expression of cell cycle genes could be mediated through the regulation of TFs. Consistent with this hypothesis, 21 TFs were among the 223 direct miRNA targets (**Figure 4D**; **Table S5A**). A striking finding was that the majority of these TFs have documented roles in cell cycle regulation, including *E2F2*, which is part of the complex controlling cell cycle progression, and many TFs known to regulate cell growth (e.g. *ZC3H10*, *ZNF783*, *ZBTB46*, *ZBTB5*, *ZFYVE26*, *TP53, EZH1, HIF3A, DPF2, TEAD3*). In addition, directly miRNA-regulated TFs included TFs involved in the regulation of beta cell development and maturation, such as *NKX6.1* (Schaffer et al., 2013; Taylor et al., 2013) and the thyroid hormone receptor *THRA*, consistent with the role of thyroid hormone in beta cell maturation (Matsuda et al., 2017). Reflective of their shared seed sequence, TFs directly regulated by Let-7g and Let-7a showed complete overlap, while miR-200a and miR-375 mostly regulated separate sets of TFs (**Figure 4E**). This analysis indicates that Let-7g, Let-7a, miR-200a, and miR-375 jointly change the transcriptional landscape in PE cells by down-regulating expression of different sets of TFs.

### miRNAs regulate a network of cell cycle genes in pancreatic progenitor cells

Having identified a set of TFs as direct miRNA targets, we next sought to determine whether these TFs could act down-stream of the miRNAs to regulate cell cycle genes. To test this, we constructed and subsequently probed a miRNA-gene regulatory network, linking the four candidate miRNAs and their direct TF targets to poly-miR-regulated genes predicted to be target genes of the TFs (**Figure 5A**; **Figure S5A,B**). First, to identify TF binding events close to poly-miR-regulated genes (down- and up-regulated), we used ATAC-seq data from PE cells and islets and mapped open chromatin regions surrounding transcriptional start sites (TSSs; closest within 10 kb) of these genes (n = 241,922 sites; FDR < 0.01, MACS2) (**Figure 5A**; **Figure S5A**; see Materials and Methods). Second, to pinpoint identified candidate TF-bound regions with likely impact on gene regulation during the PE to islet transition, we identified ATAC sites exhibiting dynamics in histone modifications between PE cells and islets. We focused on H3K4me3 and H3K27ac, two highly dynamic histone modifications during development (Wang et al., 2015; Xie et al., 2013) that have been associated with active promoters (H3K4me3) and active promoters and enhancers (H3K27ac) (Creyghton et al., 2010; Heintzman et al., 2009). We then tested whether changes in these histone marks are accompanied by expression changes of proximal genes. As predicted, an increase in H3K4me3 and H3K27ac deposition in PE compared to islets was associated with higher mRNA levels (*p* = 5.3×10^−134^; Mann Whitney test), while a decrease was associated with lower mRNA levels (*p* = 1.9×10^−36^). Finally, to construct the network, we linked open chromatin regions with dynamic histone marks to miRNA-regulated TFs by identifying those regions with a matching TF binding motif. Validating our miRNA-gene regulatory network, GO analysis showed that the 1,307 genes comprising the network were enriched for cell cycle regulators (**Figure S5C**; **Table 6**).

**Figure 5.**
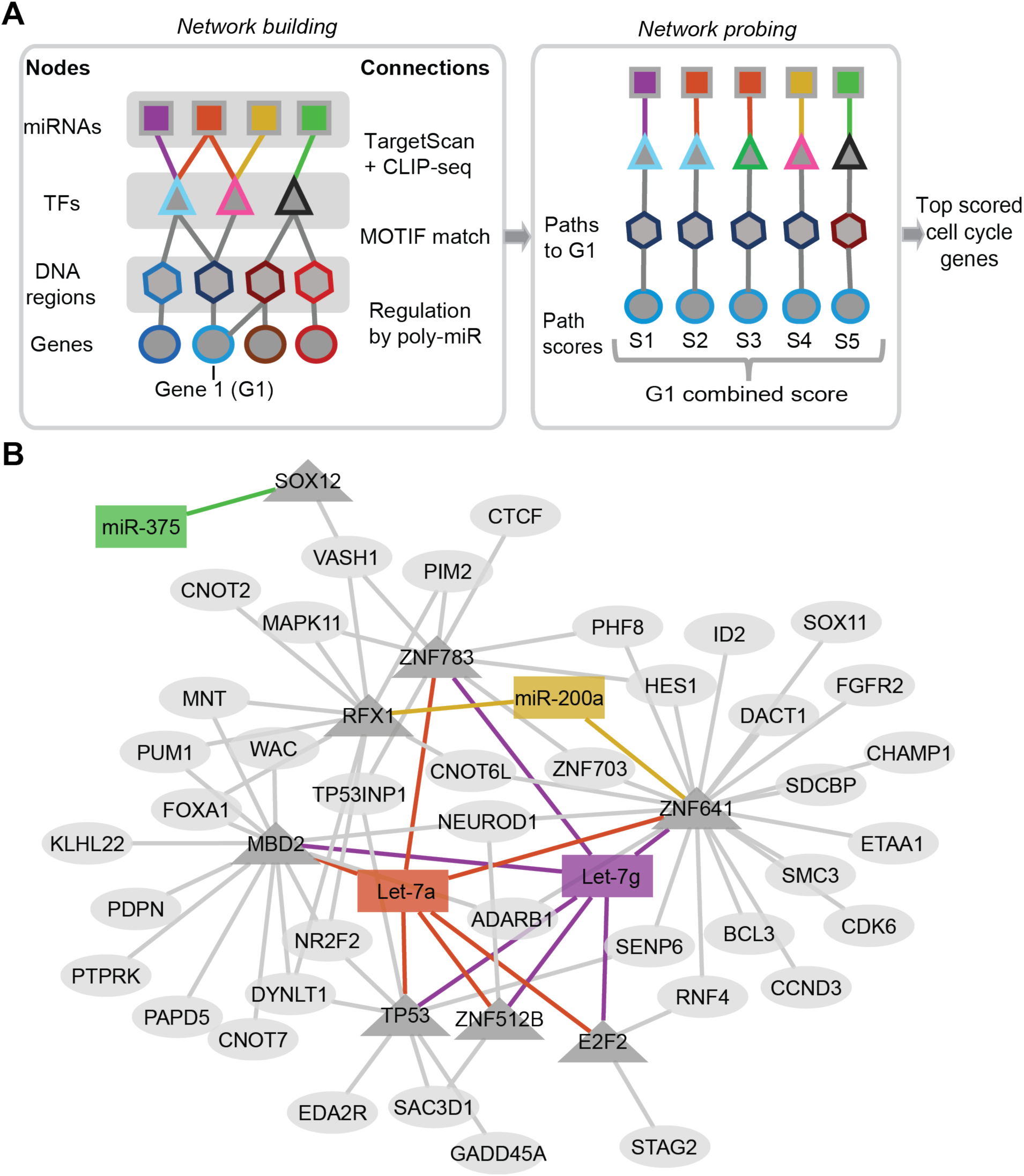
Beta cell-enriched miRNAs regulate a network of cell cycle genes. **(A)** Schematic of approach to identify core network of miRNA-regulated transcription factors and down-stream target genes. Building of network (left) and probing of network (right) is summarized. The nodes of the graph represent miRNAs [squares; Let-7g (purple), Let-7a (red), miR-200a (yellow), and miR-375 (green)], TFs (triangles), TF binding regions (hexagons) and genes (circles). **(B)** Predicted network of 40 highest scoring cell cycle genes based on network in (A) with miRNAs depicted as rectangles, TFs as triangles, and genes as ovals. TF, transcription factor; G, gene; S, score. See also Figure S5, Table S6, and Table S7.

Having validated our approach of linking putative TF binding events to changes in gene transcription during the PE to islet transition, we next assembled all data into a structured graph (**Figure 5A**; **Figure S5A**) consisting of different types of nodes that represent the individual datasets, namely the four candidate miRNAs (squares), their direct target TFs (triangles), predicted TF binding regions (hexagons), and indirect miRNA target genes (circles). Each connection between nodes (i.e. edge) was given a score representing the strength of their association, as inferred from miRNA-target databases (a), from algorithms matching TF motifs to DNA sequences (b) or from differential regulation of the connected gene (c) (see **Figure S5A** for details). A combined score was then computed for each possible path in the network from a miRNA to a gene. The score for an individual gene (G1) regulated by a miRNA is the sum of the edge scores (S1= a1 + b1 + c1). To account for synergistic effects of different miRNAs on an individual gene, we then computed a combined score representing the connectivity of the four miRNAs to G1. According to this scoring system, a higher rank is assigned to genes with strong connectivity of individual miRNA-mediated paths and characterized by a synergistic effect of more than one miRNA (**Figure S5B, Table S7**). The resulting network of the 40 highest scoring cell cycle genes demonstrates direct connectivity of the four miRNAs with cell cycle regulators through several TFs, including *SOX12, RFX1, TP53, E2F2, MBD2, ZNF512B, ZNF783,* and *ZNF641* (**Figure 5B**). Of interest is the identification of *NEUROD1* as an indirect miRNA target of Let-7 miRNAs and miR200a. Neurod1 is a TF that has been shown to induce cell cycle exit and to regulate endocrine cell differentiation in model organisms (Ahnfelt-Ronne et al., 2007; Mutoh et al., 1998). Our network analysis identifies a core network of miRNAs, TFs as their direct targets, and down-stream genes with likely roles in cell cycle regulation and endocrine cell differentiation.

### miRNAs regulate beta cell differentiation by promoting cell cycle exit

We next determined whether forced expression of Let-7g, Let-7a, miR-200a, and miR-375 represses cell cycle progression in hESC-derived pancreatic progenitor cells, as predicted by our computational analysis. We transduced PE cells with the poly-miR lentiviral construct and differentiated these cells for another six days as 3D aggregates to the early pancreatic endocrine (EN) stage, when insulin^+^ cells are first present (**Figure 6A**). Sectioned aggregates were then stained for the proliferation marker Ki-67. Validating our computational prediction, forced expression of the miRNAs reduced the percentage of Ki-67^+^ cells (**Figure 6B,C**).

**Figure 6.**
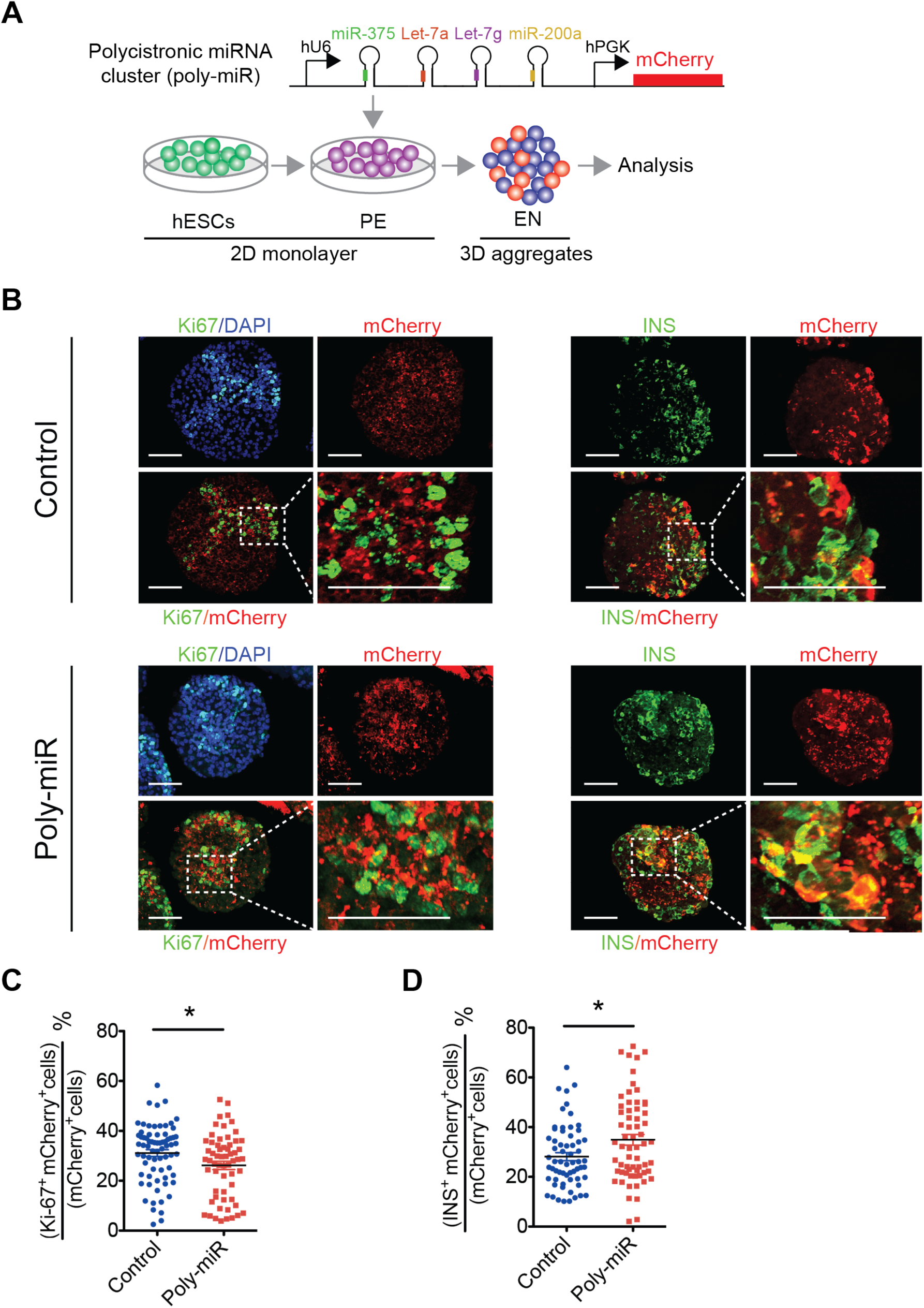
Beta cell-enriched miRNAs regulate cell cycle exit and beta cell differentiation. **(A)** Workflow to test effects of miRNAs on cell proliferation and beta cell differentiation during the transition of hESC-derived pancreatic endoderm (PE) to the early endocrine (EN) cell stage. Early PE stage cells were transduced with a lentivirus expressing a polycistronic construct for the indicated miRNAs (poly-miR) and mCherry, cultured in 2D until the end of the PE stage, aggregated, differentiated in 3D to the EN stage, sectioned, and stained for Ki-67 and insulin (INS). **(B)** Representative images showing immunofluorescence staining for Ki-67 (left) and INS (right) together with mCherry and DAPI at the EN stage for control vector (top) or poly-miR (bottom) transduced aggregates. Scale bar, 50 μm. **(C,D)** Percentage of Ki-67^+^ cells **(C)** and INS+ cells **(D)** in the mCherry+ cell population. Data are shown as mean ± S.E.M. (n = 3 biological replicates, each dot represents cell counts from one aggregate). \**p* < 0.05, Student’s t-test. See also Figure S6.

Since cell cycle exit and endocrine cell differentiation are tightly coupled (Kim et al., 2015; Miyatsuka et al., 2011; Piccand et al., 2014), we tested whether the reduction in Ki-67^+^ cells after miRNA over-expression was associated with an increase in the number of insulin^+^ cells. Indeed, we observed a higher percentage of insulin^+^ cells in aggregates expressing the poly-miR construct compared to vector-transduced aggregates (**Figure 6B,D**). The bias of our culture conditions for the differentiation of insulin^+^ cells (**Figure S6**) precluded quantification of other endocrine cell types. Taken together, our data support a model whereby endocrine-enriched Let-7g, Let-7a, miR-200a, and miR-375 are part of a gene regulatory network that triggers cell cycle exit to promote endocrine cell differentiation.

## DISCUSSION

Here, we identified 13 miRNAs that are induced during human endocrine cell differentiation. We then determined the top four candidate miRNAs with the highest predictive value of regulating endocrine cell differentiation and experimentally show that these miRNAs induce cell cycle exit of pancreatic progenitor cells. By constructing an integrated miRNA-gene regulatory network of endocrine cell differentiation, we provide evidence that these miRNAs jointly contribute to endocrine cell differentiation by regulating mRNA levels cell cycle-associated TFs.

To analyze how islet cell-enriched miRNAs cooperate to drive endocrine cell differentiation, we developed a computational method to model the relationship of miRNAs, TFs, and miRNA-regulated genes. Our computational model builds on a previously published approach for constructing miRNA regulatory networks (Gosline et al., 2016) and integrates chromatin state and expression data to build a multi-layer network. Our approach differs in several key aspects from published methodologies. First, it incorporates predictions from both CLIP-seq data and TargetScan into a combined score that is assigned to network edges. In addition, our scoring system focuses on a set of miRNAs identified experimentally and weighs the number of miRNAs contributing to each path, accounting for the synergistic effects of miRNAs on downstream gene expression changes. As such, the algorithm presented here can be applied to other cellular contexts with matching miRNA/mRNA/chromatin data and provides a useful framework for the prediction of miRNA effects.

We found that islet cell-enriched miRNAs Let-7g, Let-7a, miR-200a, and miR-375 repress different transcripts involved in cell cycle regulation and jointly drive cell cycle exit and endocrine cell differentiation. All four miRNAs have been implicated in the regulation of cell proliferation in other contexts. Like the Let-7 family miRNAs studied here, Let-7b inhibits proliferation and induces neural differentiation when overexpressed in neural progenitors (Zhao et al., 2010). Furthermore, Let-7b, miR-200a, and miR-375 have been shown to induce cell cycle arrest in tumor cells (Liu et al., 2012; Uhlmann et al., 2010; Wang et al., 2011). Likewise, acute overexpression of miR-375 in dedifferentiated beta cells reduces their proliferation (Nathan et al., 2015). This anti-proliferative effect of miR-375 is opposite to observations in *miR-375*-deficient mice, which exhibit decreased beta cell proliferation (Poy et al., 2009). Since these mice carry a germline mutation of *miR-375*, it is possible that the observed decrease in beta cell proliferation is the consequence of a developmental defect rather than a reflection of miR-375 directly regulating inhibitors of cell cycle progression.

Pancreatic endocrine cell differentiation is tightly linked to cell cycle exit. In both mice and humans, endocrine cell differentiation depends on the TF NGN3 (encoded by *NEUROG3*) (Gradwohl et al., 2000; McGrath et al., 2015), which commits pancreatic progenitors to the endocrine lineage and promotes cell cycle exit by inducing the cell cycle inhibitors Cdkn1a (p21/CIP1) and Pak3 (Miyatsuka et al., 2011; Piccand et al., 2014). We observed no effect of either combined or individual Let-7g, Let-7a, miR-200a, and miR-375 overexpression on *NEUROG3* mRNA levels (**Table S3A-E**), suggesting that these miRNAs exert their effect on proliferation independent of NGN3. However, Let-7g, Let-7a, miR-200a, and miR-375 expression with the poly-miR construct significantly induced the NGN3 target gene and endocrine differentiation factor *NeuroD1* (Ahnfelt-Ronne et al., 2007). Based on our computational model, these miRNAs are predicted to modulate *NEUROD1* expression indirectly through down-regulation of *NEUROD1* upstream TFs. Given that NEUROD1 can promote cell cycle exit through direct activation of *Cdkn1a* (Mutoh et al., 1998), miRNA-mediated modulation of *NEUROD1* mRNA levels likely has a significant contribution to the observed effects of islet-enriched miRNAs on cell proliferation and differentiation.

Gain- and loss-of-function studies in model organisms have shown that the repressive effects of miRNAs on their targets is generally modest, which has led to the view that miRNAs act to fine-tune gene expression. Consistent with this view, we observed relatively small effects of miRNA over-expression on gene expression, cell proliferation, and endocrine cell differentiation. However, these results do not mean that the miRNAs are not important for endocrine cell differentiation. We over-expressed islet cell-enriched miRNAs in an in vitro system where growth factor conditions have been optimized for efficient beta cell differentiation. Therefore, the miRNAs might not be limiting factors in this experimental context. Studies in model organisms underscore the idea that miRNAs confer robustness to developmental processes and become limiting under conditions of stress. For example, loss of miR-7 has little effect on *Drosophila* sensory organ development under normal conditions, but when environmental stresses are added to the developing organism miR-7 becomes necessary (Li et al., 2009). Similar examples exist in worms and mice, where miRNA deletions lead to significant development perturbations only on sensitized backgrounds or under stress (Brenner et al., 2010; Chivukula et al., 2014). Further illustrating that miRNAs can have significant biological effects in specific contexts, miRNAs have been shown to drastically augment reprogramming efficiencies (Anokye-Danso et al., 2011; Yoo et al., 2011). Therefore, the here-identified beta cell-enriched miRNAs could be exploited to develop reprogramming strategies to generate beta cells from other cell types.

## ACKNOWLEDGEMENTS

Human pancreatic islets were provided by the Integrated Islet Distribution Program (UC4 DK098085). We acknowledge the UCSD IGM Genomics Center for next generation sequencing (P30 DK063491). This work was supported by the National Institutes of Health (R01 DK068471 and R01 DK078803 to M.S.), the California Institute for Regenerative Medicine (RB4-06144 to M.S.) and postdoctoral fellowships from the Juvenile Diabetes Research Foundation (3-2015-83 to W.J., 3-2012-177 to A.W., and 3-2017-386 to K.V.N.N.) and the Larry L. Hillblom Foundation (2015-D-021-FEL to B.G.).

## AUTHOR CONTRIBUTIONS

W.J., F.M., and M.S. conceived the project and designed the experiments. W.J., B.G., J.W., C.Z., K.V.N.N., and H.P.S. performed experiments. F.M., Y.S., A.W., and J.C. conducted bioinformatics analysis. N.V. contributed ATAC-seq analysis, K.H.K. provided small RNA-seq data, and K.F. provided islet RNA-seq and islet ATAC-seq data. W.J, F.M., B.G., and M.S. interpreted data. W.J., F.M., A.C.C., and M.S. wrote the manuscript.

## SUPPLEMENTAL INFORMATION

**Figure S1:**
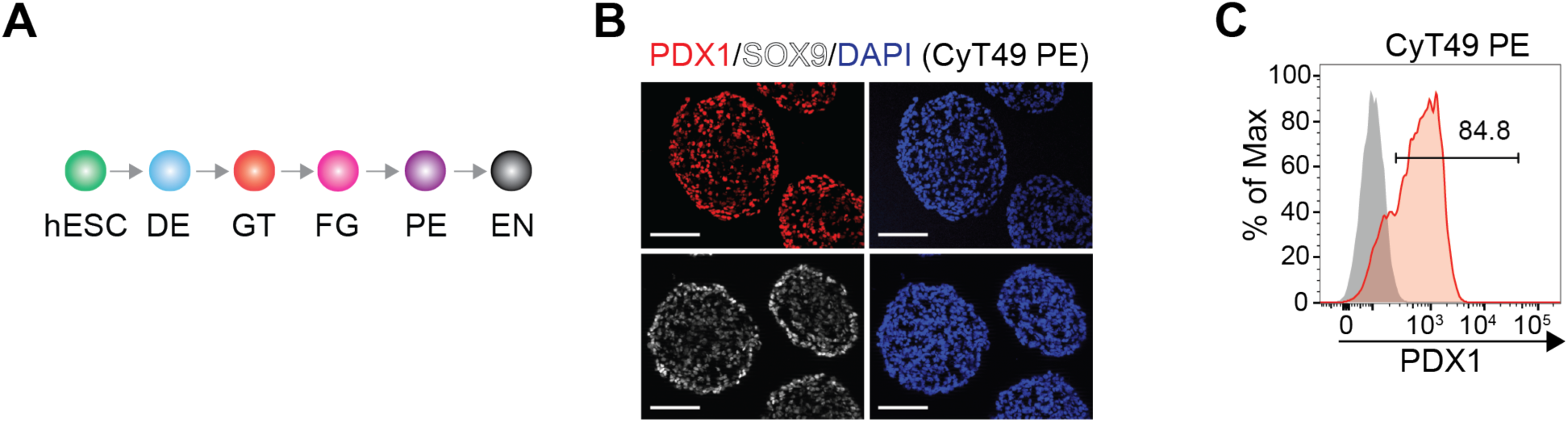
Pancreatic endoderm differentiated from CyT49 hESCs. Related to Figure 1. **(A)** Schematic of the hESC-based differentiation strategy. **(B)** Immunofluorescence staining of PE cell aggregate sections for PE-specific markers PDX1 and SOX9. Scale bar, 50 μm. **(C)** Representative flow cytometry analysis at PE stage for PDX1. hESC, human embryonic stem cells; DE, definitive endoderm; GT, gut tube; FG, posterior foregut; PE, pancreatic endoderm; EN, endocrine cell stage.

**Figure S2.**
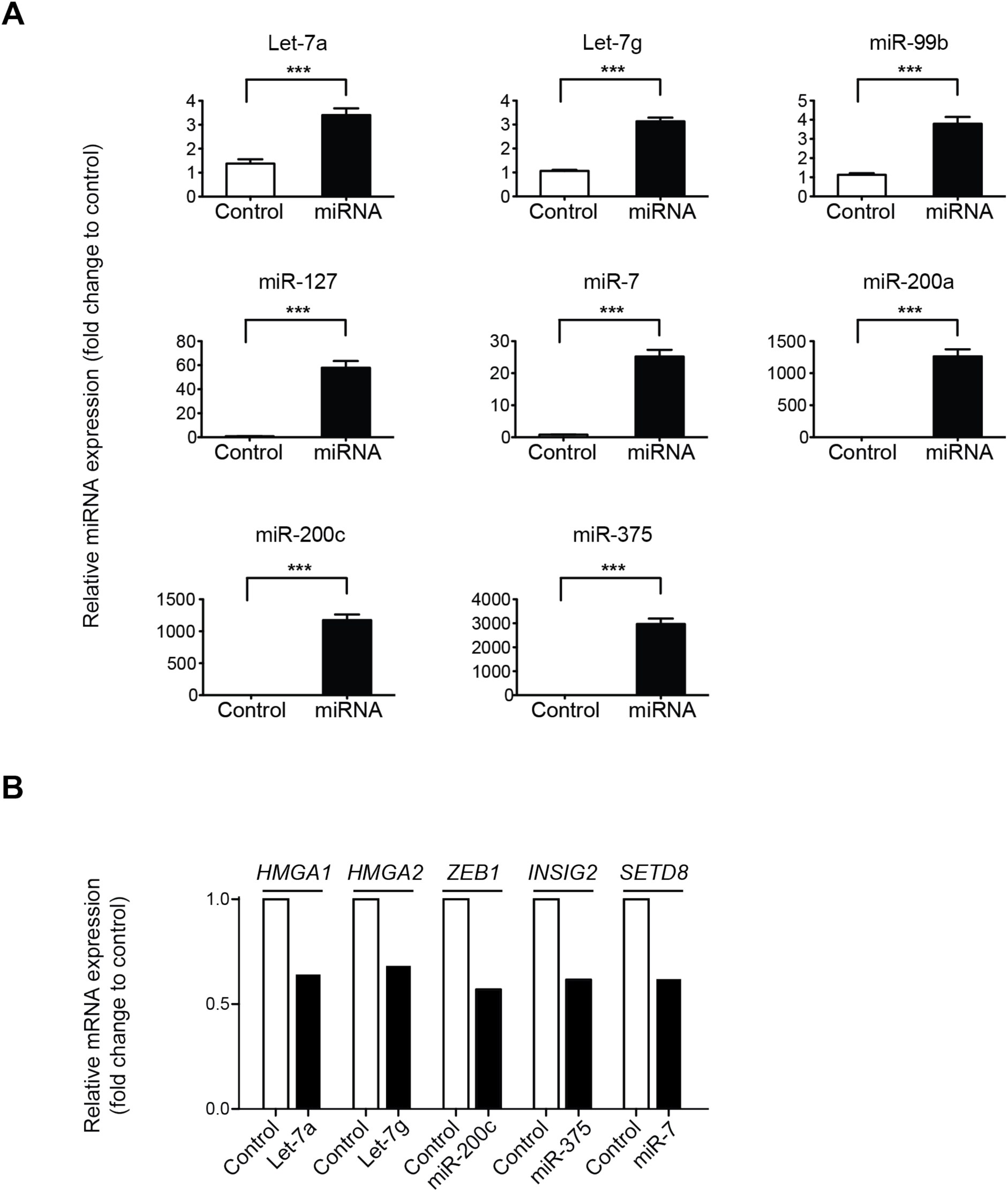
Forced expression of individual miRNAs in HeLa cells. Related to Figure 2. **(A)** Relative expression of indicated miRNAs determined by Taqman qPCR in HeLa cells 24 h after transfection with miRNAs or vector control. Data are shown as mean ± S.E.M. (n = 3 biological replicates). \*\*\**p* < 0.001; Student’s t-test. **(B)** Expression of previously published target mRNAs for indicated miRNAs determined by RNA-seq analysis of HeLa cells transfected with miRNAs or vector control (n = 1).

**Figure S3.**
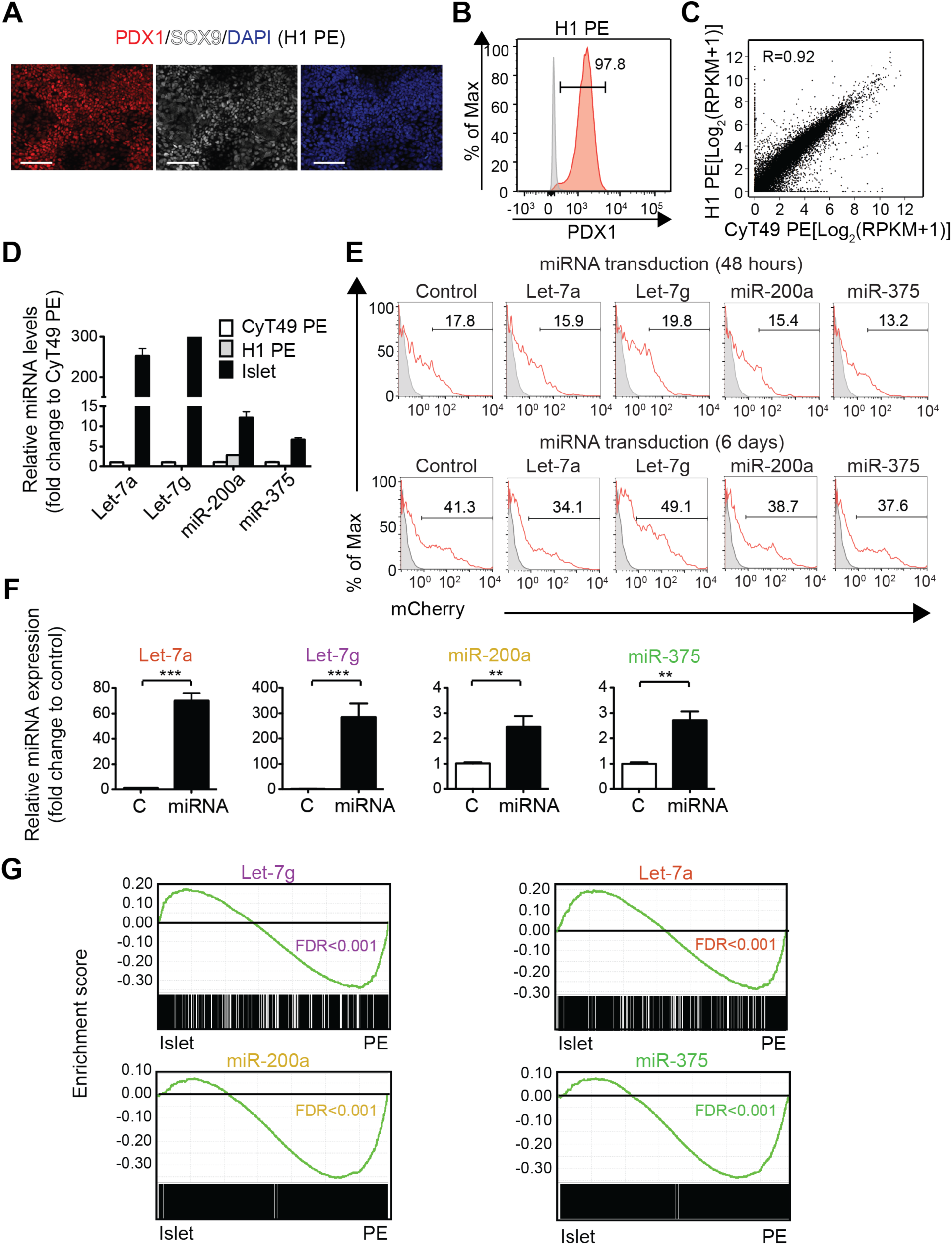
Forced expression of individual miRNAs in hESC-derived pancreatic progenitor cells. Related to Figure 3. **(A)** Immunofluorescence staining of pancreatic endoderm (PE) differentiated from H1 hESCs for PDX1 and SOX9. Scale bar, 50 μm. **(B)** Representative flow cytometry analysis at PE stage for PDX1. **(C)** Scatter plot showing correlation in mRNA expression between PE cells derived from H1 hESCs and CyT49 hESCs. **(D)** Expression of indicated miRNAs in H1-derived PE cells and islets relative to CyT49-derived PE cells determined by Taqman qPCR. Data are shown as mean ± S.E.M. (n = 3 technical replicates). **(E)** Representative flow cytometry analysis for mCherry 48 h (top row) and 6 days (bottom row) after lentiviral transduction with miRNA-mCherry constructs. Gating for cell sorting is shown. **(F)** Relative expression of indicated miRNAs determined by Taqman qPCR in H1 PE cells 48 h after lentiviral transduction with miRNAs or vector control (C). Data are shown as mean ± S.E.M. (n = 3 biological replicates). \*\**p* < 0.01, \*\*\**p* < 0.001; Student’s t-test. **(G)** GSEA plots showing enrichment of genes repressed by Let7g, Let7a, miR-200a, and miR-375 in islets (n = 3) compared to PE (n =2). False Discovery Rate (FDR) is shown.

**Figure S4:**
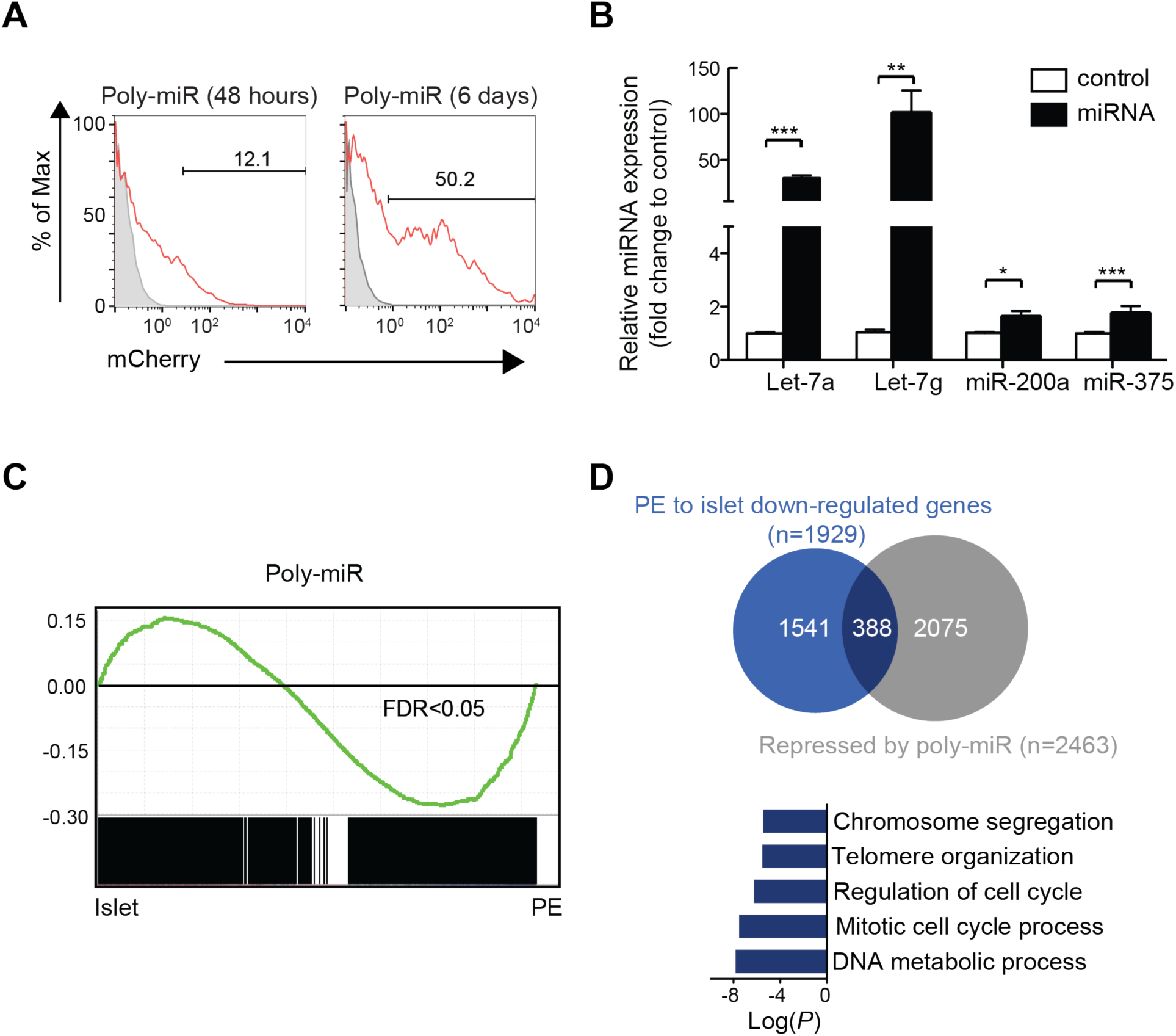
Forced expression of a polycistronic construct for four miRNAs in hESC-derived pancreatic progenitor cells. Related to Figure 4. **(A)** Representative flow cytometry analysis for mCherry 48 h (left) and 6 days (right) after lentiviral transduction with poly-miR-mCherry construct. Gating for cell sorting is shown. **(B)** Relative expression of indicated miRNAs determined by Taqman qPCR in H1 PE cells 48 h after lentiviral transduction with a vector control or a polycistronic construct containing Let-7g, Let-7a, miR-200a, and miR-375 (poly-miR). Data are shown as mean ± S.E.M. (n = 3 biological replicates). \*\**P* < 0.01, \*\*\**P* < 0.001; Student’s t-test. **(C)** GSEA plot showing enrichment of genes repressed by the poly-miR construct in islets compared to PE. False Discovery Rate (FDR) is shown. **(D)** Venn diagram showing the overlap between genes down-regulated in islets compared to PE (blue) and genes repressed by the poly-miR construct (grey). Top five GO categories enriched among genes repressed by the poly-miR construct and down-regulated in islets compared to PE are shown on the bottom.

**Figure S5:**
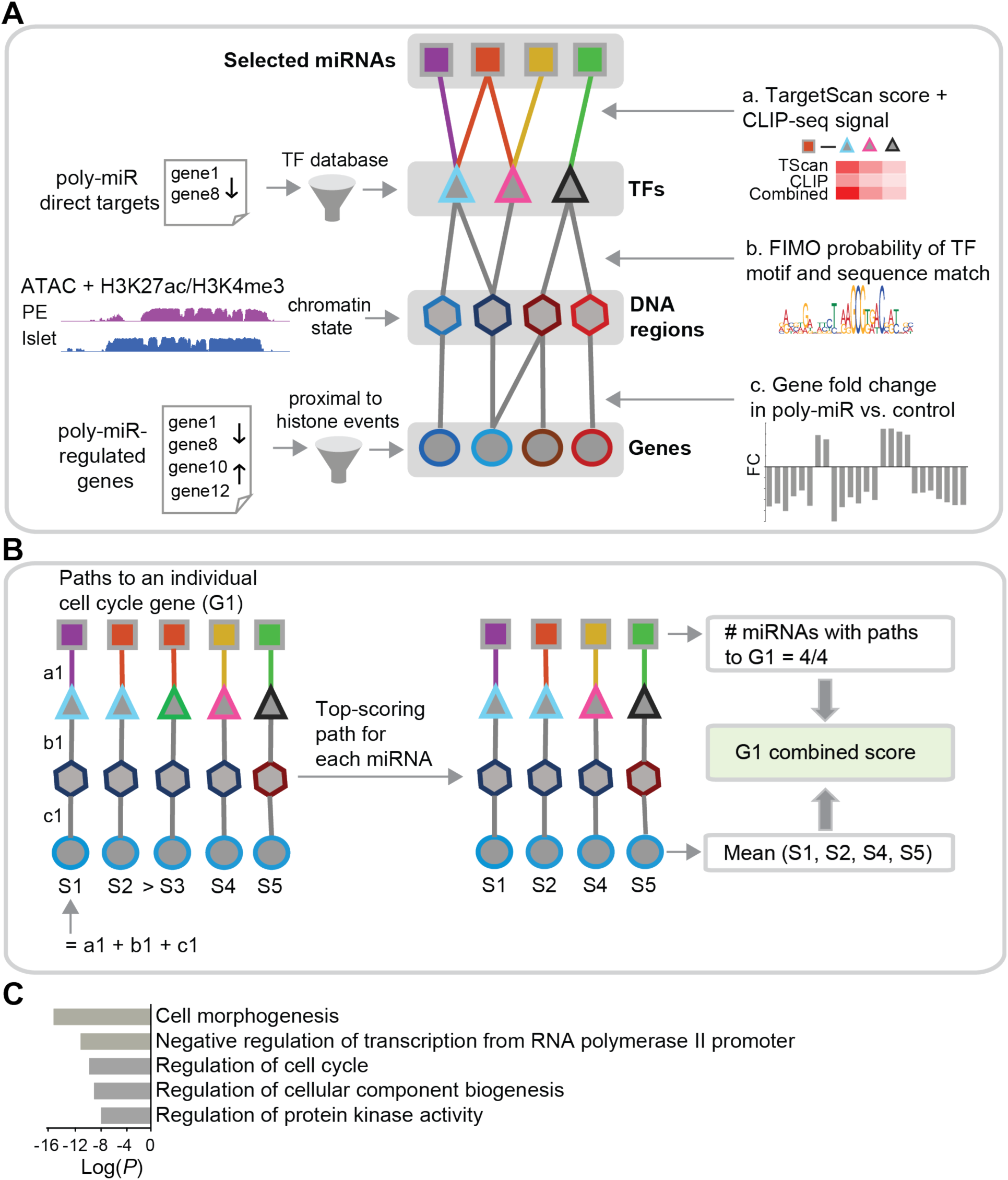
Approach to identify core network of miRNA-regulated genes in pancreatic progenitor cells. Related to Figure 5. **(A,B)** Schematic of approach to identify core network of miRNA-regulated transcription factors and cell cycle genes. Building of network **(A)** and probing of network **(B)** is shown. The nodes of the graph represent miRNAs [squares; Let-7g (purple), Let-7a (red), miR-200a (yellow), and miR-375 (green)], transcription factors (triangles), transcription factor binding regions (hexagons) and genes (circles). In **(A)**, the source for each node is indicated on the left, while evidence (indicated by a, b, and c) used to calculate scores is indicated on the right. In **(B)**, path-based scoring of an individual gene [e.g. gene 1 (G1)] is shown. See Materials and Methods for details. **(C)** Top five GO categories enriched in genes comprising the miRNA-regulated core network in (A). TF, transcription factor; G, gene; S, score.

**Figure S6:**
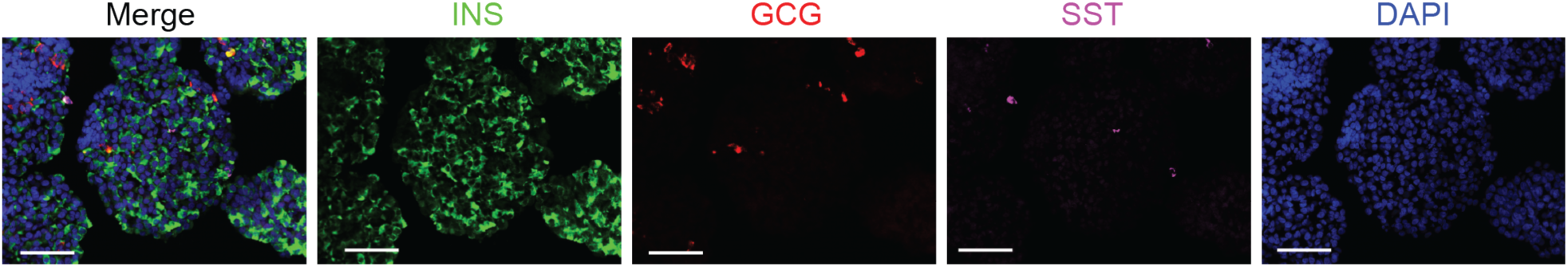
Endocrine cell types differentiated from H1 hESCs. Related to Figure 6. Immunofluorescence staining of sections from endocrine cell stage aggregates for insulin (INS), glucagon (GCG), and somatostatin (SST). Scale bar, 50 μm.

### Supplemental Tables

**Table S1. miRNA expression in PE, pancreatic alpha and beta cells.**

**(A)** miRNA expression values (RPM) in PE, pancreatic alpha and beta cells.

**(B)** miRNA expression fold-change in beta cells vs PE.

**(C)** miRNA expression fold-change in alpha cells vs PE. Table S1 is associated with Figure 1.

**Table S2. Genes regulated by miRNAs in Hela cells.**

**(A)** Let-7a, **(B)** Let-7g, **(C)** miR-99b, **(D)** miR-127, **(E)** miR-124, **(F)** miR-7, **(G)** miR-200a, **(H)** miR-200c, and **(I)** miR-375 regulated genes. Table S2 is associated with Figure 2.

**Table S3. Genes regulated by individual miRNA and poly-miR in PE cells.**

**(A)** Let-7a, **(B)** Let-7g, **(C)** miR-200a, **(D)** miR-375, and **(E)** poly-miR regulated genes. Table S3 is associated with Figures 3 and 4.

**Table S4. GO analysis of individual miRNA and poly-miR repressed genes.**

GO analysis of genes that decrease in expression in islet compared to PE and are repressed by **(A)** Let-7a, **(B)** Let-7g, **(C)** miR-200a, **(D)** miR-375, and **(E)** poly-miR. Table S4 is associated with Figures 3 and 4.

**Table S5. Analysis of direct targets for poly-miR.**

**(A)** List of poly-miR direct targets. **(B)** GO analysis of poly-miR direct targets. Table S5 is associated with Figure 4.

**Table S6. GO analysis of all genes in the miRNA-regulated network.** Table S6 is associated with Figure 5.

**Table S7. Network-based ranking of cell cycle genes.** Table S7 is associated with Figure 5.

## STAR METHODS

### CONTACT FOR REAGENT AND RESOURCE SHARING

Further information and requests for reagents may be directed and will be fulfilled by the corresponding author Maike Sander.

### EXPERIMENTAL MODEL AND SUBJECT DETAILS

#### Human islets

Human pancreatic islets were obtained from three human cadaveric organ donors through the Integrated Islet Distribution Program (IIDP). Human pancreatic islets had ≥ 90% purity and ≥ 90% viability. Upon receipt, islets were handpicked and immediately processed for RNA extraction.

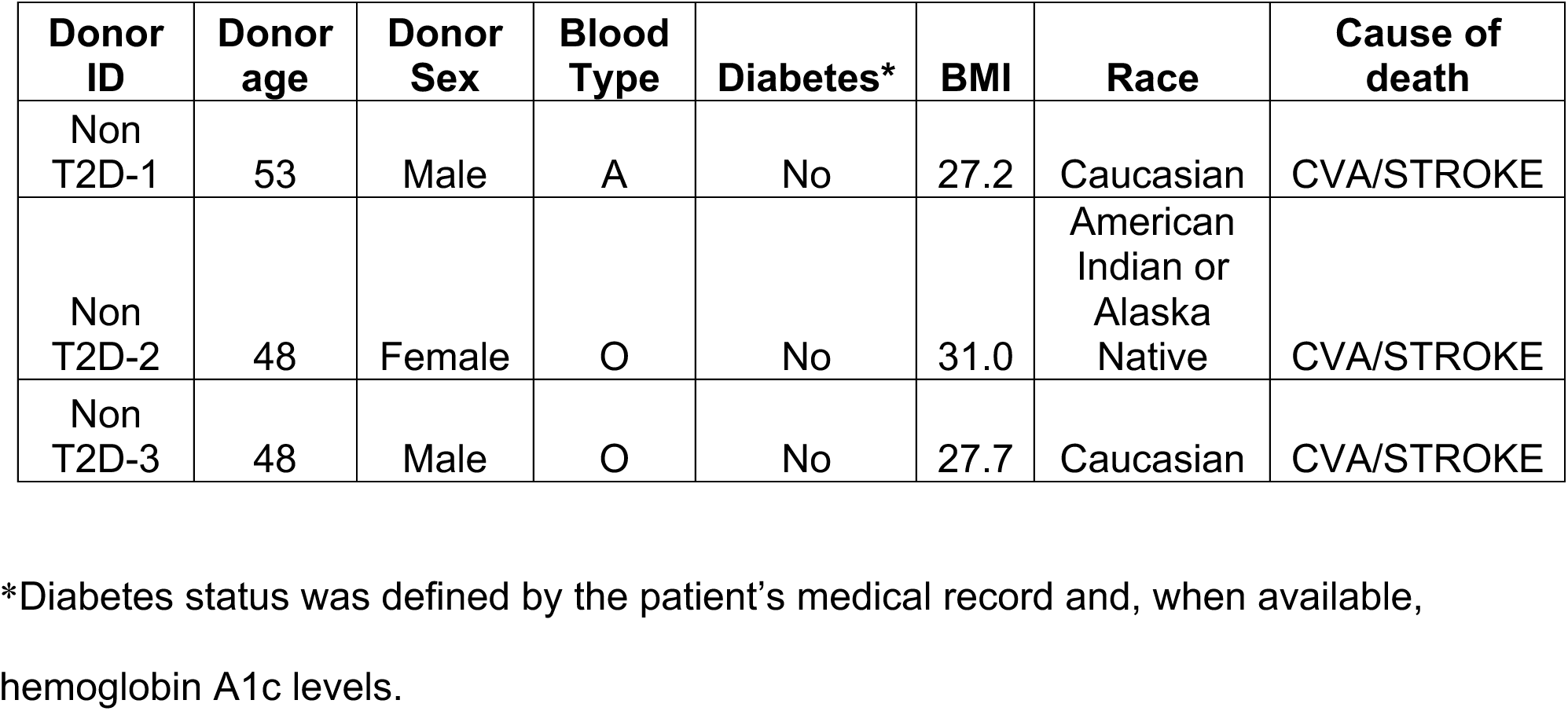

#### Maintenance and differentiation of H1 hESCs

hESC research was approved by the University of California, San Diego, Institutional Review Board and Embryonic Stem Cell Research Oversight Committee. All hESC experiments were performed in H1 hESCs with the exception of miRNA expression profiling, for which CyT49 hESCs were used.

H1 hESCs were maintained and differentiated as described with some modifications (Rezania et al., 2014). In brief, hESCs were cultured in mTeSR1 media (Stem Cell Technologies) and propagated by passaging cells every 3 to 4 days using Accutase (eBioscience) for enzymatic cell dissociation. For differentiation of H1 cells, we employed a 2D monolayer culture format up to day 11 of differentiation. Cells were then dissociated using accutase for 10 min, reaggregated by plating the cells in a low attachment 6-well plate on an orbital shaker (100 rpm) in a 37 °C incubator. Cells were subsequently cultured in suspension from day 11-14.

On day 0, dissociated hESCs were resuspended in mTeSR1 media (see media compositions below) and seeded onto Matrigel-coated 12-well plates by adding 1 ml of cell suspension (∼8 x 10^5^ cells/well) to each well. The following day, undifferentiated cells were washed in stage 1 medium and then differentiated using a multi-step protocol with stage-specific media (see below) and daily media changes.

All stage-specific base media were comprised of MCDB 131 medium (Thermo Fisher Scientific) supplemented with NaHCO_3_, GlutaMAX, D-Glucose, and BSA using the following concentrations:

Stage 1/2 medium: MCDB 131 medium, 1.5 g/L NaHCO3, 1X GlutaMAX, 10 mM D-Glucose, 0.5% BSA

Stage 3/4 medium: MCDB 131 medium, 2.5 g/L NaHCO3, 1X GlutaMAX, 10 mM D-glucose, 2% BSA

Stage 5 medium: MCDB 131 medium, 1.5 g/L NaHCO3, 1X GlutaMAX, 20 mM D-glucose, 2% BSA

Media compositions for each stage were as follows:

Stage 1 (day 0-2): base medium, 100 ng/ml Activin A, 25 ng/ml Wnt3a (day 0). Day 1-2: base medium, 100 ng/ml Activin A

Stage 2 (day 3-5): base medium, 0.25 mM L-Ascorbic Acid (Vitamin C), 50 ng/mL FGF7

Stage 3 (day 6-7): base medium, 0.25 mM L-Ascorbic Acid, 50 ng/mL FGF7, 0.25 μM SANT-1, 1 μM Retinoic Acid, 100 nM LDN193189, 1:200 ITS-X, 200 nM TPB

Stage 4 (day 8-10): base medium, 0.25 mM L-Ascorbic Acid, 2 ng/mL FGF7, 0.25 μM SANT-1, 0.1 μM Retinoic Acid, 200 nM LDN193189, 1:200 ITS-X, 100nM TPB

Stage 5 (day 11-14): base medium, 0.25 μM SANT-1, 0.05 μM Retinoic Acid, 100 nM LDN193189, 1:200 ITS-X, 1 μM T3, 10 μM ALK5 inhibitor II, 10 μM ZnSO4, and 10 μg/mL Heparin, 10 μM ROCK inhibitor.

End of stage 1 = definitive endoderm

End of stage 2 = gut tube

End of stage 3 = posterior foregut

End of stage 4 = pancreatic endoderm

End of stage 5 = endocrine cells

#### Maintenance and differentiation of CyT49 hESCs

CyT49 hESCs were maintained and differentiated as described (Xie et al., 2013). Propagation of CyT49 hESCs was carried out by passing cells every 3 to 4 days using Accutase™ (eBioscience) for enzymatic cell dissociation, and with 10% (v/v) human AB serum (Valley Biomedical) included in the hESC media the day of passage. hESCs were seeded into tissue culture flasks at a density of 50,000 cells/cm2.

CyT49 hESC media was comprised of DMEM/F12 (Corning; 45000-346) supplemented with 10% (v/v) KnockOut™ Serum Replacement (Thermo Fisher Scientific), 1X MEM non-essential amino acids (Thermo Fisher Scientific), 1X GlutaMAX™ (Thermo Fisher Scientific), 1% (v/v) penicillin-streptomycin (Thermo Fisher Scientific), 0.1mM 2-mercaptoethanol (Thermo Fisher Scientific), 10ng/mL Activin A (R&D Systems), and 10ng/mL Heregulin-β1 (PeproTech).

Pancreatic differentiation of CyT49 hESCs was performed as previously described (Schulz et al., 2012). Briefly, we employed a suspension-based format using rotational culture. Undifferentiated hESCs were aggregated by preparing a single cell suspension in hESC media at 1 × 106 cells/mL and overnight culture in six-well ultra-low attachment plates (Costar) with 5.5ml per well on an orbital rotator (Innova2000, New Brunswick Scientific) at 95 rpm. The following day, undifferentiated aggregates were washed in RPMI media (Corning) and then differentiated using a multistep protocol with daily media changes and continued orbital rotation at either 95 rpm or at 105 rpm on days 4 to 8.

Stage 1/2 medium: RPMI medium (Corning), 0.2 % (vol/vol) FBS, 1X GlutaMAX

Stage 3/4 medium: DMEM High Glucose medium (HyClone), 0.5X B-27 Supplement, 1X GlutaMAX

Media compositions for each stage were as follows:

Stage 1 (Day 0-1): Day 0: RPMI/FBS, 100ng/mL Activin A, 50ng/mL mouse Wnt3a, 1:5000 ITS. Day 1: RPMI/FBS, 100 ng/mL Activin A, 1:5000 ITS.

Stage 2 (Day 2-4): Day 2: RPMI/FBS, 2.5μM TGFβ R1 kinase inhibitor IV, 25ng/mL KGF, 1:1000 ITS. Day 3-4: RPMI/FBS, 25ng/mL KGF, 1:1000 ITS

Stage 3 (Day 5-7): DMEM/B27, 3nM TTNPB, 0.25mM KAAD-Cyclopamine, 50ng/mL Noggin Stage 4 (Day 7-10): DMEM/B27, 50ng/mL KGF, 50ng/mL EGF.

End of stage 1 = definitive endoderm

End of stage 2 = gut tube

End of stage 3 = posterior foregut

End of stage 4 = pancreatic endoderm

#### Cell lines

HEK293T and HeLa cells were maintained in DMEM containing 100 units/mL penicillin and 100 mg/mL streptomycin sulfate supplemented with 10% fetal bovine serum (FBS).

### METHOD DETAILS

#### Immunocytochemistry

Cells were washed twice before fixation with 4% paraformaldehyde in PBS for either 30 min at room temperature, or overnight at 4°C. Cells were then washed three times with PBS and incubated in 30% sucrose at 4°C overnight before mounting in Optimal Cutting Temperature Compound (Tissue-Tek) and sectioning at 10 μm. Immunocytochemistry was performed as described (Xie et al., 2013). The following primary antibodies and dilutions were used: guinea pig anti-PDX1 (gift from Dr. Christopher Wright, Vanderbilt University) 1:1000; rabbit anti-SOX9 (Millipore, AB5535) 1:1000; rabbit anti-Ki-67 (ThermoFisher, RM-9106-S1) 1:200; guinea pig anti-insulin (LINCO, 4011-01) 1:1000. Secondary antibodies were Cy5-, Cy3-, or Alex488-conjugated donkey antibodies against guinea pig or rabbit (Jackson Immuno Research Laboratories). Images were acquired on a Zeiss Axio-Observer-Z1 microscope with a Zeiss AxioCam digital camera and figures prepared with Adobe Photoshop CS6/Illustrator CS5.

To determine the percentage of Ki-67^+^ cells in the mCherry^+^ cell population, at least ten sections from different aggregates were analyzed per hPSC differentiation. For each condition, three independent hPSC differentiations were performed. Ki-67^+^ and mCherry^+^ cells were quantified using HALO software (PerkinElmer Inc).

#### Fluorescence-activated cell sorting and intracellular flow cytometry

hESC-derived PE cells were dissociated to a single-cell suspension with Accutase (Stemcell Technologies) at 37°C for 10 min. Accutase was neutralized with FACS sorting buffer [1% (wt/vol) FBS, 1 mM EDTA, 25mM Hepes, PBS]. FACS was performed on a FACS Fortessa equipped with FACS DiVa software (BD Biosciences). Cells were sorted into Trizol for RNA analysis. For intracellular flow cytometry, dissociated cells were fixed, permeabilized with BD Cytoperm/Cytofix (BD Bioscience), and stained with anti-PDX1-PE conjugated antibody (BD Biosciences, 562161; 1:20) at room temperature for 30 min, washed, and resuspended in FACS buffer. Flow cytometry analysis was performed on FACSCanto II (BD Biosciences) and analyzed with FlowJo software (FloJo LLC).

#### TaqMan microRNA assay

qRT-PCR for miRNAs was performed using TaqMan MicroRNA Reverse Transcription Kit (Applied Biosystems, Cat. No. 4366596). Briefly, 10 ng of total RNA was reverse transcribed using RT primers from the TaqMan MicroRNA Assay kit [Applied Biosystems; probe catalogue numbers: Let-7a (000377), Let-7g (002282), miR-127 (000452), miR-200a (000502), miR-375 (000564), miR-7 (000268), miR-99b (000436), and RUN44 (001094). qRT-PCR was performed on a Bio-rad CFX96 real-time system using the TaqMan Universal PCR Master Mix (Applied Biosystems, Cat. No. 4324018) and TaqMan probes from the TaqMan MicroRNA Assay kit. miRNA levels were determined on three independent samples and values were normalized to endogenous snoRNA RNU44.

#### miRNA expression vector construction and transfection

To generate miRNA expression vectors, 270 nt of the miRNA gene primary transcript, including the 22 nt mature miRNA and 125 nt of genomic sequence flanking each side of the miRNA (Chen et al., 2004), were amplified and placed under the control of the human U6 pol III promoter in the pLKO.3G backbone (Addgene, plasmid #14748). Let-7a and Let-7g were expressed with mutations in their loop sequence to block LIN28 binding and ensure proper miRNA processing (Piskounova et al., 2008). HeLa cells were transfected using a 1 mg/ml PEI solution (Polysciences) to achieve >95% transfection efficiency. Cells were harvested and RNA extracted 24 hr after transfection.

For the polycistronic miRNA expression vector, a gBlock gene fragment encompassing mIR-375, let-7a, let-7g, and mIR-200a was cloned into a modified version of pLKO.3G, in which GFP was exchanged for mCherry,

#### Lentivirus production and transduction of PE cells

High-titer lentiviral supernatants were generated by co-transfection of the miRNA expression vector and the lentiviral packaging construct into HEK293T cells as described (Xie et al., 2013). Briefly, miRNA expression vectors were cotransfected with the pCMV-R8.74 (Addgene, #22036) and pMD2.G (Addgene, #12259) expression plasmids into HEK293T cells using a 1mg/ml PEI solution (Polysciences). Lentiviral supernatants were collected at 48 hr and 72 hr after transfection. Lentiviruses were concentrated by ultracentrifugation for 120 min at 19,500 rpm using a Beckman SW28 ultracentrifuge rotor at 4°C. The titer routinely achieved was 5*10^8^∼10^9^ TU/ml. For PE cell transductions, H1 hESCs were differentiated to the PE stage (day 8 of differentiation) in monolayer cultures and transduced with lentivirus at a MOI of 2. For RNA analysis, cells were collected 48 hr after transduction.

#### Small RNA sequencing and data analysis

Small RNA-seq data from sorted human alpha and beta cells have been described (Kameswaran et al., 2014). RNA from PE stage CyT49 hESC cultures was isolated using the miRVana miRNA Isolation kit (Thermo Fisher Scientific). 3 μg of RNA was used for library preparation using the TruSeq Small RNA sample preparation kit (Illumina) and a Pippin Prep (Sage Science) for size selection with a 3% cassette (CSD3010). RNA was prepared for sequencing using the Illumina protocol (Illumina FC-102-1009) and amplified libraries were sequenced on an Illumina Genome Analyzer II (Illumina FC-104-1003). Sequenced libraries were processed as described (Kameswaran et al., 2014). miRNAs with sample values below 1 RPM were excluded from the analysis. There was one replicate each for hESC-derived PE, alpha, and beta cells. Each miRNA expression value was log_2_-transformed and displayed in a heatmap.

#### RNA sequencing, mapping and data analysis

RNA quality was assessed using TapeStation (Agilent Technologies). Libraries were prepared according to the instructions of Illumina’s TruSeq RNA library prep kit. Libraries were quantified using High Sensitivity DNA screen tape (Agilent Technologies) and Qubit dsDNA High Sensitivity (Life Technologies) assays. Finally, libraries were multiplexed and sequenced on a HiSeq 2500 (Illumina) sequencer using single-end sequencing.

RNA-seq samples were mapped to the UCSC human transcriptome (hg19/GRCh37) by the Spliced Transcripts Alignment to a Reference (STAR) aligner (STAR-STAR_2.4.0f1), allowing for up to 10 mismatches (Dobin et al., 2013). Only reads aligned uniquely to one genomic location were retained for subsequent analysis. Expression levels of all genes were quantified by Cufflink (https://github.com/cole-trapnell-lab/cufflinks) using only reads with exact matches. Genes with average RPKM above 1 were retained for further analyses.

Differentially expressed genes were identified using a permutation test, with the number of permutations set to 1000. Briefly, all the samples were shuffled, fold changes were computed to obtain a null distribution, and a *P-*value was estimated for each gene’s fold change as a cumulative probability from the null distribution. For comparison of PE and islet data, batch effects were removed using ComBat (Johnson et al., 2007).

#### Gene Set Enrichment Analysis (GSEA) and Gene Ontology (GO) analysis

We applied GSEA (http://www.broad.mit.edu/gsea), which scores a-priori defined gene sets in two different conditions (Subramanian et al., 2005). GSEA (http://www.broad.mit.edu/gsea) was run with the number of permutations for *P-*value computation set to 1000. We used genes significantly repressed by miRNAs (*P* < 0.05, permutation test) as gene sets to determine coordinated regulation in islets compared to PE samples. Gene sets with a false discovery rate of < 0.05 were considered significantly enriched.

Enrichment of gene sets for Gene Ontology (GO) terms was tested using Metascape (Tripathi et al., 2015).

#### ChIP-seq and ATAC-seq data analysis

ChIP-seq and ATAC-seq reads were mapped to the human genome (hg19/GRCh37) using Bowtie (Langmead et al., 2009) and BWA (Li and Durbin, 2009), respectively, and visualized using the UCSC Genome Browser (Kent et al., 2002). Unmapped reads were discarded. After mapping, SAMtools (Li et al., 2009) was used to remove duplicate sequences and merge samples. Here, “SAMtools view -Sbq 30” was used to filter out reads with mapping quality less than 30, “SAMtools rmdup” was used to remove duplicated reads, and “SAMtools merge” was used to merge files of the same histone marker or input condition. ChIP-seq and ATAC-seq analysis was performed in two biological replicates for PE and 4-5 donors for islet. The Pearson correlation among biological replicates ranged from 64% to 96% for human islets and 91% to 96% for PE.

HOMER (Heinz et al., 2010) as used to call ChIP-seq peaks using “findPeaks function” with “–style histone” to call peaks. Stage- and condition-matched input DNA controls were used as background when calling peaks. MACS2 (Zhang et al., 2008) was used to call peaks from ATAC-seq data, with parameters “shift set to 100 bps, smoothing window of 200 bps” and with “nolambda” and “nomodel” flags on.

To link changes in chromatin to gene expression changes, we first defined differential H3K27ac and H3K4me3 peaks in PE and islet (adjusted *P*<0.05, “getDifferentialPeaksReplicates” function in HOMER) and then used BEDtools (Quinlan and Hall, 2010) to identify overlapping ATAC peaks in PE or islet using a ± 1.5 kb window from the summit of the ATAC peak. Next, we identified the nearest TSS within a 10kb window of the H3K27ac or H3K4me3 peak. We then assessed the concordance of the directionality of changes in gene expression and histone marks by evaluating whether genes near regions showing gain or loss of H3K27ac or H3K4me3 in PE versus islet exhibit significant concordant expression changes (Mann-Whitney test).

#### Network building

CLIP-seq signal, mRNA expression, ATAC-seq and ChIP-seq binding data were encoded in a graphical model depicted in Figures 5 and Supplemental Figure S5 by adapting a previously published algorithm (Gosline et al., 2016). Nodes arranged in four different layers, corresponding to miRNAs, TFs, DNA regions, and genes, were identified and connected as follows: For each of the four selected miRNAs, predicted target genes were retrieved through the TargetScan repository (http://www.targetscan.org/vert_71/). For the pairs of miRNA-target gene identified, the corresponding CLIP-seq signal was collected from previously published data (Kameswaran et al., 2014). Among the targets, TFs in the second layer of the network were selected based on down-regulation by poly-miR transfection compared to control with *P* < 0.05 and with an annotation in the TF Animal Database (Zhang et al., 2015). In the third layer of the network, we selected DNA regions showing significant changes in PE versus islet for H3K27ac or H3K4me3 (see previous paragraph) with the nearest gene showing down-regulation by poly-miR transfection, hereafter referred to as *Rsel*. The *Rsel* DNA regions were filtered for links to the selected TFs by scoring the match of their binding motifs with the DNA regions in the network. Briefly, motifs of selected TFs were extracted from a collection of databases, including JASPAR (http://jaspar.genereg.net/cgi-bin/jaspar_db.pl), Homococo (http://hocomoco11.autosome.ru/), and ENCODE-related data sets (Aylward et al., 2018) and scored for matches with narrow regions spanning 300bp around the peak summits of each *Rsel*. Log-odds scores and corresponding *P*-values were obtained using the MEME Suite tool FIMO (http://meme-suite.org/tools/fimo) with default parameters. The last layer was defined by considering genes proximal to DNA regions, as described above, and filtering for those differentially regulated in poly-miR versus control (*P* < 0.05).

A score representing the strength of the association was computed for each pair of connected nodes in the different network layers, as follows: Given the TargetScan context++ score *S_i_* (Agarwal et al., 2015) and the CLIP-seq signal *S_j_* of each miRNA-TF association, their values were normalized in a 0-1 range and combined as *a* = (1 − *S_i_*)(1 − *S_j_*) (Szklarczyk et al., 2015). Scores of edges connecting TFs to *Rsel* regions were defined as the *b* = 1 − *q*, *q* being the q-value returned by FIMO, representing the probability of obtaining the log-odds ratio scores of the matches by chance. Scores from DNA regions to genes were defined as the absolute value of the Log2 Fold Change of each gene *x* in poly-miR versus control data: *c* = |*Log*2*FC*(*x*)| A combined score was computed for each possible path in the network starting from a miRNA to a gene, by adding the contribution of the different layers, as: *S* = *a* + *b* + *c*.

#### Network-based gene ranking

Given an individual gene, G1, all network paths connecting a miRNA to G1 were considered with their corresponding scores and compared for gene ranking as follows: First, among the paths connecting the same miRNA to G1, only the one with the highest score was retained, obtaining a score S_i_ for each of the *k* miRNAs showing an indirect link to G1, k<=N, with N equal to the number of miRNAs in the network (N=4). A first score summarizing the strength of the association of these retrieved network paths was computed as *S_net_*(*G*1) = *mean*(*S_i_*), *i* ranging from one to *k*. A second score, accounting for the synergistic effect of several miRNAs on the same gene, was computed as proportion of miRNAs in at least one network path linking to the selected gene G1: *S_mir_*(*G*1) = *k*/*N*. Finally, a combined score for G1 was obtained as a weighted sum of *S_net_* and *S_mir_*, i.e. *S_comb_* (*G*_1_) = *w* ∗ *S_net_* (*G*1) + *w* ∗ *S_mir_* (*G*1), with *w* set to 0.5. This procedure was applied to score individual genes annotated to cell cycle regulation in the Gene Ontology (GO:0051726) and genes were ranked based on *S_comb_* values.

### QUANTIFICATION AND STATISTICAL ANALYSIS

Statistical parameters including FDR, R, and *p* values are reported in the Figures and the Figure Legends.

### DATA AVAILABILITY

#### Accession numbers

All RNA-seq and ATAC-seq data generated in this study can be found at GEO with accession number GSE115327 (reviewer code: inivicayhrgzzyp).

Accession numbers for additional data used in this study are as follows: GSE52314 (small RNA-seq, sorted alpha and beta cells); GSE51924 (CLIP-seq, human islets); E-MTAB-1086 (RNA-seq, PE cells); GSE54471 (H3K27ac, H3K4me3 ChIP-seq, PE cells); GSE51311, E-MTAB-1919, E-MTAB-189, and E-MTAB-191 (H3K27ac, H3K4me3 ChIP-seq, human islets).

RNA-seq and ATAC-seq data for human islets can be accessed through: https://www.t2depigenome.org/publications/4eacc3f8-428c-4271-881c-d18e3b423fdf/ (User name; Password: dgareviewer).

